# Cell-based Simulations of Biased Epithelial Lung Growth

**DOI:** 10.1101/747188

**Authors:** Anna Stopka, Marco Kokic, Dagmar Iber

## Abstract

During morphogenesis, epithelial tubes elongate. In case of the mammalian lung, biased elongation has been linked to a bias in cell shape and cell division, but it has remained unclear whether a bias in cell shape along the axis of outgrowth is sufficient for biased outgrowth and how it arises. Here, we use our 2D cell-based tissue simulation software `LBIBCell` to investigate the conditions for biased epithelial outgrowth. We show that the observed bias in cell shape and cell division can result in the observed bias in outgrowth only in case of strong cortical tension, and comparison to biological data suggests that the cortical tension in epithelia is likely sufficient. We explore mechanisms that may result in the observed bias in cell division and cell shapes. To this end, we test the possibility that the surrounding tissue or extracellular matrix acts as a mechanical constraint that biases growth in longitudinal direction. While external compressive forces can result in the observed bias in outgrowth, we find that they do not result in the observed bias in cell shapes. We conclude that other mechanisms must exist that generate the bias in lung epithelial outgrowth.

## 1. Introduction

Epithelial tubes are an essential component of many organs, and their development has been studied in many different model organisms. In *Drosophila*, where overall growth is limited by the exoskeleton, tube elongation is mostly associated with changes in cell shape (Figure 1a) and cell position (Figure 1b), which can also include the recruitment of surrounding cells [1]. For example, in the *Drosophila* salivary gland and trachea, tubes elongate by cell shape change in combination with cell migration. In the *Drosophila* tracheal system, dynamic filopodia and lamellipodia have been observed in response to Fgf signaling, which would support directed cell migration [2]. Mammalian organs grow extensively during development, and it is largely unknown how the outgrowth can become biased [1]. In this article, we focus on the biased elongation of epithelial tubes during mouse lung development.

**Figure 1.**
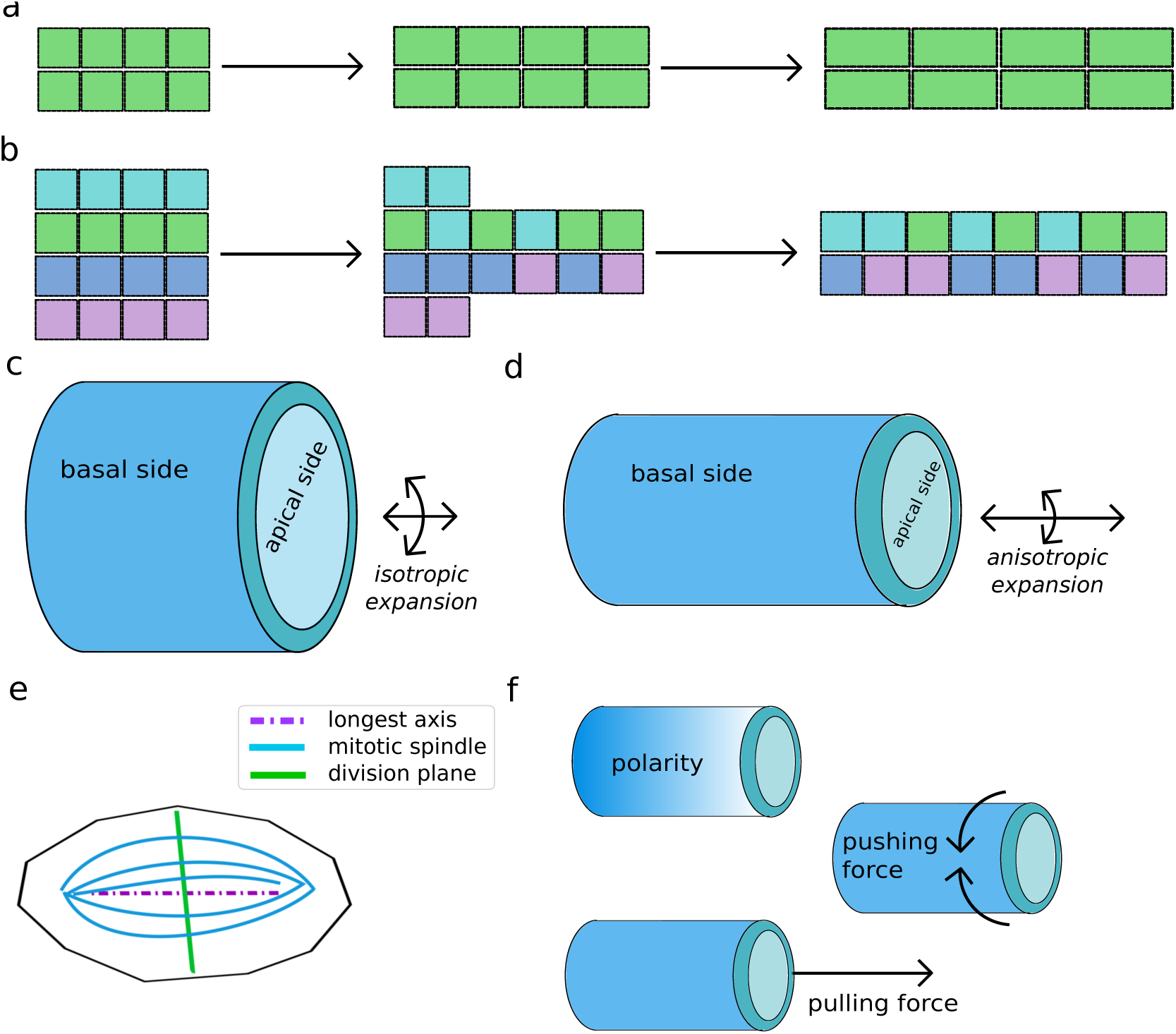
Mechanisms controlling tube elongation. (a-b) Tube elongation via cell shape changes and cell rearrangements. (c-d) An epithelial tube consisting of a monolayer with an outer basal side and an inner apical side facing the lumen. The tube expands either isotropically, i.e. it grows by the same factor in length and circumference, or the tube expands anisotropically, e.g. it grows more in length than in circumference. (e) A cell dividing according to Hertwig’s rule. (f) Three possible mechanisms to achieve anisotropic tube expansion: polarity along the tube e.g. via signaling gradients, compressive forces along the circumference, or pulling forces along the length of the tube.

The developing mouse lung offers examples of both, isotropic (Figure 1c) and anisotropic (Figure 1d) tube expansion. Thus, the mouse embryonic tracheal epithelium expands isotropically, i.e. the circumference widens to the same extent as the tube elongates [3], whereas the mouse embryonic lung bronchi show biased outgrowth, i.e. the epithelial tube lengthens more than it widens [4, 5]. The biased outgrowth of embryonic mouse lung bronchi has been related to a bias in the orientation of the mitotic spindles of the dividing cells: *∼* 42% of the dividing cells orient their mitotic spindle in the direction of elongation [4]. Similarly, in the mouse kidney, a bias in the orientation of the mitotic spindles of dividing cells is observed during renal tube elongation [6]. According to Hertwig’s rule, cells divide through their mass point, and perpendicular to their longest axis (Figure 1e) [7, 8]. Indeed, the bias in cell division is accompanied by a bias in cell shape [5]. But how does the bias in cell shape arise in the embryonic lung?

The planar cell polarity (PCP) pathway plays an important role in regulating the mitotic spindle angle distribution [9, 10]. Loss of the protocadherin Fat4 disrupts PCP signaling, oriented cell division, and biased outgrowth in the mouse kidney, and additional heterozygosity in Van Gogh-like 2 (Vangl2), a component of the PCP signaling pathway, leads to an even stronger manifestation of the phenotype [6]. Whether the PCP pathway is also responsible for the bias in cell divisions in the mouse embryonic lung is unclear as Vangl2*^Lp^* mice have normal airway shape and spindle orientation [4]. However, independent of whether or not the PCP pathway is involved, it does not address the question of how the direction of outgrowth is defined in the first place.

In principle, such a bias could originate from a polarity along the tube, from a pulling force at the tip, or from a mechanical constraint that limits circumferential expansion (Figure 1f) [11]. The most obvious polarity cue is provided by fibroblastic growth factor (Fgf), a well-known chemoattractant for lung buds [12]. However, while lung buds grow towards a source of Fgf in culture, there is no evidence that Fgf signaling gradients exist along outgrowing buds, and bud outgrowth is observed when the epithelium is exposed to uniform Fgf concentrations [13]. In fact, computational modeling suggests that a ligand-receptor based Turing mechanisms concentrates Fgf signaling in spots where new branches grow out [14, 15], and thus also contradicts the formation of a gradient along the epithelial tube.

Alternatively, Fgf signaling could mediate elongating outgrowth by generating a pulling force at the tip. In fact, Fgf2 signaling has been shown to enhance motility and persistence of epithelial cells at the tip of epithelial tubes in the mammary gland [16]. A pulling force would act on the cells behind the tip via the cell-cell adhesion bonds, and could thus generate a bias in cell shape along the entire tube. In case of pulling forces at the tip, actin-rich protrusions would be expected at the tip. However, in the mammary gland, leading cell extensions or specialized actin-rich protrusions, such as filopodia and lamellipodia, are not observed at the front of advancing mammary ducts [17]. Instead, F-actin is enriched along lateral cell surfaces and colocalized with Zonula occludens-1 (ZO-1), a tight junction-associated protein, at luminal surfaces [17].

As an alternative to cell migration, biased outgrowth in the mammary gland has been linked to a mechanical constraint posed by the myoepithelium [18]. Such a mechanical constraint in circumferential direction would result in growth-dependent uniform expansion in longitudinal direction, and would thus explain the measured linear gradient in cell speeds that is observed along the elongating mammary gland tube [16]. The lung mesenchyme could, in principle, play a similar role in the embryonic lung though smooth muscles mature only at later stages and are not as tightly associated as in a myoepithelium [19]. In the developing kidney, however, smooth muscles are largely absent, and branches elongate also in the absence of mesenchyme [20]. Finally, in plants hoop stress has been suggested to induce an anisotropy in the cell walls [21], such that growth is limited in circumferential direction. While the cytoplasmic hydrostatic pressure and extracellular matrix (ECM) stiffness are orders of magnitude smaller than in plants, a bias in ECM stiffness could, in principle, limit expansion in circumferential direction also in mammalian organs.

An additional challenge in explaining epithelial outgrowth is posed by the visco-elastic nature of tissue. While cortical tension together with cell-cell adhesion interactions result in an elastic tissue behavior, cellular remodeling, cell migration, cell shape changes and rearrangements [22, 23] result in a fluid-like behavior [24, 25]. Whether the observed bias in cell division is sufficient to explain the bias in outgrowth will depend on the fluidity of the tissue, and on potential external forces that may bias the growth process. In this context, it is interesting to note that the bias in cell division and in epithelial outgrowth is lost in embryonic lungs that express a constitutive active form of KRas (*Shh^cre/^*^+^; *KRas^LSL/^*^+^) [4], which activates Erk, a target of Fgf signaling [26]. KRas signaling in airway epithelial cells has previously been linked to changes in cell shape and motility by affecting cortical actin [27]. Expression of a constitutive active form of KRas (KRas*^G^*^12^*^D^*) in pancreatic ducts results in a redistribution of phosphorylated myosin light chain 2 (pMLC2) inside the cell, so that its concentration declines on the apical side and increases on the basal side [28]. Constitutive active KRas may thus reduce cortical tension on the apical side of the epithelium, where all cell divisions occur [29].

In cases where tissue behavior emerges from processes at the single cell level, cell-based simulations can be helpful to identify the underlying mechanisms [30]. We sought to use cell-based tissue simulations to explore whether the observed bias in cell division, is, in principle, sufficient to drive the observed bias in outgrowth when taking the visco-elastic tissue properties into account, and whether constricting forces from the surrounding tissue can, in principle, explain the observed bias in cell division, cell shape, and outgrowth. We have previously developed the 2D cell-based simulation environment `LBIBCell` to simulate visco-elastic tissue dynamics [31]. `LBIBCell` models the fluid-structure interaction in a biological tissue by combining the Lattice Boltzmann (LB) [32] and Immersed Boundary (IB) [33] methods. Cells are represented as finely resolved polygons according to the IB method. The cell boundaries are elastic and cells interact with their neighbors via spring-like cell-cell junctions. The fluid behavior inside and outside of the cells is described by the LB equations. Cell-based simulations are particularly useful when quantitative data is available to inform cell-based rather than continuous models [34, 30]. In case of epithelia, quantifications of the apical polygonal lattice are available for a wide range of epithelia from *Drosophila* and mouse [4, 35, 36, 5, 37]. These can be used to adjust the mechanical parameters in the model [37, 38]. The additional biological parameters, i.e. the growth rate, the rate of apoptosis and cell division, and the orientation of the cell division axis have all been measured in embryonic mouse lungs. Using `LBIBCell`, we find that the bias in the cell division axis is sufficient to explain the observed bias in outgrowth only for high cortical tension. A mechanical constraint from the surrounding tissue, as has been proposed for the mammary gland [18], can give rise to biased outgrowth, but fails to explain the observed bias in cell shape and thus cell division orientation. We conclude that the bias in embryonic lung outgrowth can arise from the observed bias in cell division, but the bias in cell shape cannot arise from mechanical constraints imposed by the surrounding tissue. Other mechanisms must thus drive biased epithelial outgrowth of mouse lung tubes.

## 2. Methods

We use `LBIBCell` to simulate the epithelial tissue dynamics [31]. `LBIBCell` is an open-source C++ simulation environment that enables the detailed representation of 2D biological tissues at cellular resolution. As mitosis and cytokinesis are confined to the most apical epithelial region [29], our simulations focus on the apical cell dynamics (Figure 2a). We have previously described the application of `LBIBCell` to the simulation of the apical cell dynamics in epithelial tissues [37]; in the following, we provide a brief summary.

**Figure 2.**
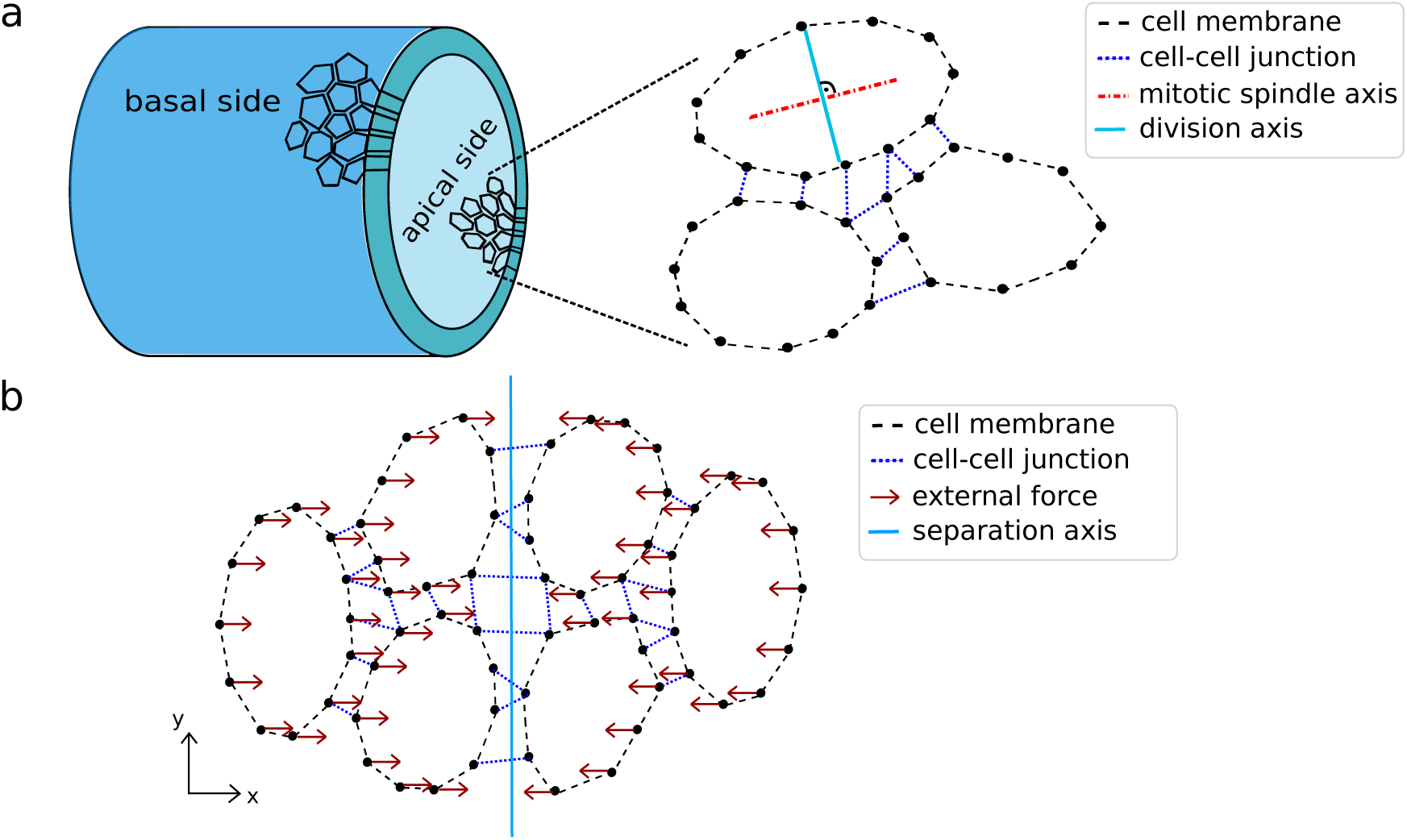
Schematic representations of simulated cells in LBIBCell. (a) *An epithelial tube consisting of a monolayer with an outer basal side and an inner apical side facing the lumen. Cells in `LBIBCell` represent the apical side of a tube. Cell cortical (membrane) tension and cell-cell junctions are represented by Hookean springs. The cell division angle is assumed to be perpendicular to the mitotic spindle axis, here the longest axis of a cell.* (b) *Illustration of the application of external forces. The simulated domain is split into two halves (blue line) and the forces are applied at each cell node acting in opposing directions*.

### Setup of LBIBCell simulations

In `LBIBCell`, every biological process is represented by means of a `BioSolver (BS)`. We used the existing BS for cell growth, cell division, apoptosis, cortical (membrane) tension, and cell-cell adhesion [31, 37], and developed a new BS to include external forces as described below. Growth of the cells is simulated by adding mass to intracellular grid points. Cells that exceed a certain cell area are divided. The cell division threshold follows a Gaussian distribution with mean *µ*_div_ and standard deviation *σ*_div_. The cell division plane was set either to follow a measured distribution (see below), or followed Hertwig’s rule [7, 8], i.e. cells were divided perpendicular to their longest axis (Figure 2a). Cells that fall below a certain cell area or that come too close to the boundaries of the simulated domain are removed. Cortical tension (called membrane tension in `LBIBCell`) and cell-cell adhesion are represented by Hookean spring forces between the cell membrane points (Figure 2a).

The BS involve a total of 14 free parameters (Table 1). Seven of these parameters affect numerical aspects, and these were set for maximal computational efficiency while avoiding numerical artefacts, as described before [37]. For the remaining seven parameters that describe biological processes we have previously defined parameter ranges that result in epithelia-like lattices [37], and we further set parameter values to recapitulate measurements in the mouse embryonic lung. Thus, the mass added to intracellular grid points per time step is severely limited by numerical stability. We used a growth source *g* = 10*^−^*^4^ and adjusted the overall simulation time to match the data. The lung tubes expand 3.6-fold between somite stage 38 to 47, which corresponds to about 25 h = 90 000 s [39], corresponding to a growth rate of 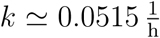, assuming exponential growth [4]. With a growth source *g* = 10 we require 36000 iteration steps to obtain a similar domain growth in the simulations, such that 2.5 s *≡* 1 LB_time_. The cell division threshold was set such that the average cell area is 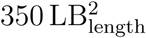 in the simulations. With a measured average cell area of 30 µm^2^ [4], the length conversion between LB units and SI units is then 0.293 µm *≡* 1 LB_length_. For numerical reasons, we set the minimal cell size, below which cells were removed (threshold for cell apoptosis and removal) to 20 LB_length_2. The cell-cell junction length was set to 0.1 LB_length_ = 30 nm, consistent with the measured E-cadherin length [40]. The only parameters, for which we do not have data from the embryonic mouse lung, are the spring constants. Accordingly, we studied their impact in parameter screens within a previously determined parameter range that results in epithelia-like tissues [37]. If not stated otherwise, we set 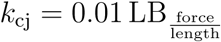 and 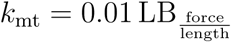.

**Table 1.** BioSolver and their parameters for the LBIBCell simulations. The `BioSolver` and their parameters are listed. When the parameter values are set within their valid range, the simulated tissues are epithelial-like.

| BioSolver | Parameter description | Dimension | Valid range [LB] | Value |
| --- | --- | --- | --- | --- |
| **Cell Growth** | mass added to intracellular grid point<br>call frequency | Mass/Time<br>(Time)<sup>-1</sup> | $\{10^{-3}, \dots, 10^{-4}\}$<br>$\geq 10^{-2}$ | $10^{-4}$<br>1 |
| **Cell Membrane Tension** | Hookean spring constant<br>call frequency | Force/Length<br>(Time)<sup>-1</sup> | $\{10^{-3}, \dots, 10^{-2}\}$<br>$\geq 10^{-2}$ | screening<br>$10^{-1}$ |
| **Cell-Cell Junction** | Hookean spring constant<br>length of intercellular cell junction<br>radius for junction building<br>call frequency | Force/Length<br>Length<br>Length<br>(Time)<sup>-1</sup> | $\{10^{-3}, \dots, 10^{-1}\}$<br>$\{10^{-2}, \dots, 10^0\}$<br>$\{0.5, \dots, 1.5\}$<br>$\geq 10^{-2}$ | screening<br>$10^{-1}$<br>1.0<br>1 ($10^{-1}$) |
| **Cell Division** | mean cell area ($\mu_{\text{div}}, \sigma_{\text{div}}$)<br>call frequency | (Length)<sup>2</sup><br>(Time)<sup>-1</sup> | $\geq 100$<br>$\leq 10^{-2}$ | $\simeq 350$<br>$10^{-2}$ |
| **Cell Apoptosis** | minimal cell area<br>call frequency | (Length)<sup>2</sup><br>(Time)<sup>-1</sup> | $\leq 50$<br>$\leq 10^{-2}$ | 20<br>$10^{-2}$ |
| **External force** | force per cell node<br>call frequency | Force<br>(Time)<sup>-1</sup> | $\{10^{-6}, \dots, 10^{-5}\}$<br>$\geq 10^{-2}$ | screening<br>1 |

### Application of external forces

To simulate the impact of external forces on tissue outgrowth, we implemented a new `BS` in `LBIBCell` that applies external forces to the cells, the `External Forces BS`. In addition to the membrane tension and the cell-cell junctions, a free force can be applied to the cell nodes, whose strength and direction are defined by a two dimensional vector. The application of this force can be coupled to cell properties such as cell type or the position of a cell. With the new `BS`, we split the simulated domain in two halves and apply forces in opposing directions to imitate the effect of a surrounding tissue that prevents the epithelial tube to expand in circumferential direction (Figure 2b).

### Cell division according to a predetermined distribution

To study the impact of biased cell division orientation on tissue growth, we developed a `Cell Division BS` that applies cell division according to a predetermined distribution. In previous versions of `LBIBCell`, dividing cells split along an individual cell division plane whose direction is either chosen based on a random number generator or perpendicular to the longest axis. In the `Cell Division BS`, the mitotic spindle angle of a cell is drawn from a given mitotic spindle angle distribution. The cell division angle is perpendicular to the mitotic spindle angle [37].

## 3. Results

### 3.1. A bias in mitotic spindle orientation results in biased outgrowth only for strong cortical tension

We sought to analyze whether the observed bias in cell shape and cell division orientation (Figure 3a-c) would be sufficient to explain the measured bias in embryonic lung tube outgrowth [4, 5]. To this end, we simulated tissue growth using parameter values that are consistent with measurements in mouse lung epithelia (Table 1), and we divided cells either according to the biased mitotic spindle angle distribution that has been experimentally measured in wildtype mouse lungs (Figure 3b) or according to the uniform mitotic spindle angle distribution of *Shh^cre/^*^+^; *Kras^LSL/^*^+^ mutant (Figure 3c) mouse lung epithelia [4]. In both cases, the simulation started with an initial tissue structure which consisted of about 100 cells, had approximately the form of a square and was positioned in the middle of the simulated domain. In case of the mitotic angle distribution from the wildtype, the final tissue adopted a rectangular tissue shape with longer sides in the respectively biased division direction (Figure 3d). On the other hand, for the experimentally measured mitotic spindle angle distribution found in *Shh^cre/^*^+^; *Kras^LSL/^*^+^ mutant lung epithelia [4], the tissue grew over time, but preserved its square shape (Figure 3e). This demonstrates that the biased mitotic spindle angle orientation in the wildtype can result in biased tissue outgrowth.

**Figure 3.**
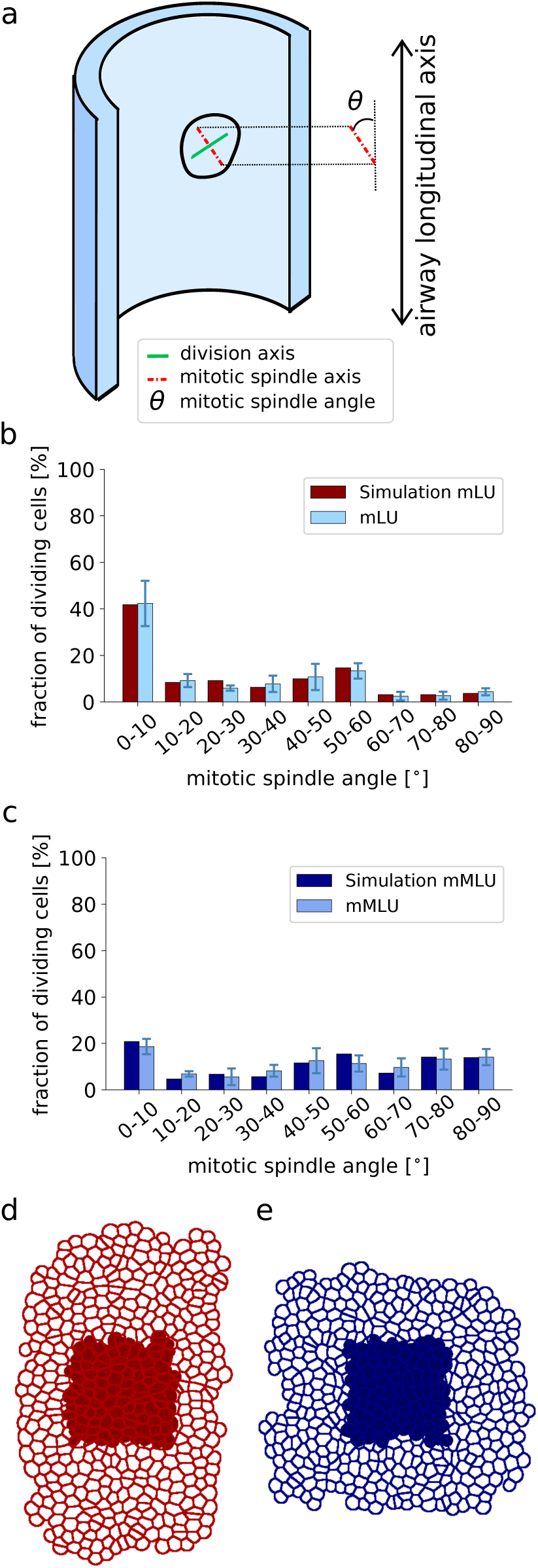
Oriented mitotic spindle angle distribution leads to biased tissue outgrowth. (a) *Schematic representation of the mitotic spindle orientation, θ, relative to the longitudinal axis of a lung tube*. (b,c) *Experimentally measured distribution of mitotic spindle orientations, θ, from (b) wildtype mouse lungs (mLU) and (c) mutant mouse lungs (mMLU)* [4] *and an exemplary simulated distribution for each case. The mean and standard deviation of are shown. (d,e) Comparison of the initial tissue configuration (filled cells) with final tissue shape of the simulation, shows* (d) *anisotropic tissue expansion in the direction of the mitotic spindle bias* (b), *and* (e) *isotropic tissue expansion in case of the mutant spindle angle distribution* (c).

We next sought to investigate the parameter dependency of biased outgrowth. Following [4], we define the bias in outgrowth as 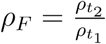, with 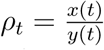; *y*(*t*) denotes the length of the tissue along the elongation axis, and *x*(*t*) the tissue length in orthogonal direction. Accordingly, the bias in outgrowth along the elongation axis (y axis) is larger, the smaller *ρ_F_*. There are only seven parameters with a biological interpretation, and these determine the growth rate, the cell area division threshold, the critical cell area below which cells delaminate or undergo apoptosis, the length of intercellular junctions, as well as the spring constants for cell-cell adhesion and cortical tension. When changing any of the non-mechanical parameters by *±*20%, the relative changes in *ρ_F_* is less than 12% (Figure 4b). In particular, the length of intercellular cell-cell junctions and the critical cell area under which cells delaminate or undergo apoptosis have only a very small impact on the bias of outgrowth, i.e. the change in *ρ_F_* is less than 5%.

**Figure 4.**
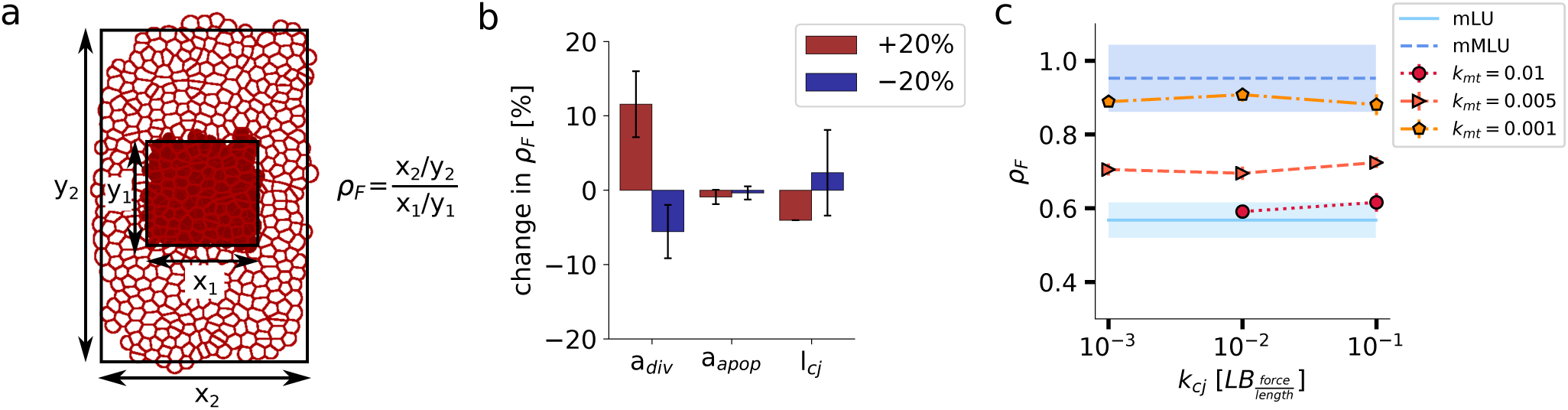
Parameter Sensitivity Analysis with regard to bias in tissue outgrowth. (a) *To determine the bias in outgrowth, the irregularly shaped tissue is approximated by a rectangle of the same area, using a least squares fit. The bias of outgrowth is then calculated by comparing the ratio between the axes at two consecutive time points. The smaller ρ_F_, the stronger is the outgrowth along the elongation axis (y axis)*. (b) *The relative changes in the airway shape change ρ_F_ for the wildtype were < 12% when changing the division area threshold, the apoptosis area and the length of cell-cell junctions by ±20%. The mean and the standard deviation are shown for each parameter set*. (c) *The airway shape change ρ_F_ depends only on cortical (membrane) tension, but not on the cell-cell junction force. The simulated outgrowth ratios were compared to the measured values for wildtype* (mLU) *and mutant* (mMLU) *mouse lung epithelia* [4].

We next screened the two mechanical parameters, the spring constants for cell-cell adhesion, *k*_cj_, and cortical (membrane) tension, *k*_mt_, within a range that gives rise to epithelial-like tissue (Table 1). We note that only a very small range of cell-cell adhesion strengths, *k*_cj_, yield an epithelial-like tissue: for lower values of the cell-cell adhesion constant, *k*_cj_, cells detach, while for higher values, cells become angularly shaped. Within this range, the cell-cell adhesion constant *k*_cj_ does not affect the bias in outgrowth (Figure 4c). On the contrary, the bias in outgrowth *ρ_F_* is highly sensitive to the cortical tension, *k*_mt_, and the outgrowth bias that is observed in lung epithelial tubes (*ρ_F_ ∼* 0.6, Figure 4c, shaded in light blue) is obtained only for the highest possible cortical (membrane) tension tension constant *k*_mt_ = 0.01.

Oriented divisions lead to biased outgrowth only for high cortical tension because of the importance of cell rounding. Upon cell division, the daughter cells are first elliptically shaped with the axis of cell division marking the long axis (Figure 5a). At high cortical tension, cells seek to round up. Cell rounding drives a redistribution of cell mass in the direction of the mitotic spindle axis, thus resulting in biased outgrowth (Figure 5a). For increasing cell size this effect is weaker (Figure 4b and Figure 5b). The effect is visible also in a tissue context (Figure 5c,d). At high cortical tension, the cells round up, and the rounding results in mass being directed towards the axis of elongation (Figure 5c). For low cortical tension, this cell rounding does not happen (Figure 5d), such that the tissue expands isotropically even though cells divide with a bias in the direction of elongation. Importantly, in all cases cells remain clonal, i.e. daughter cells remain in a close neighborhood.

**Figure 5.**
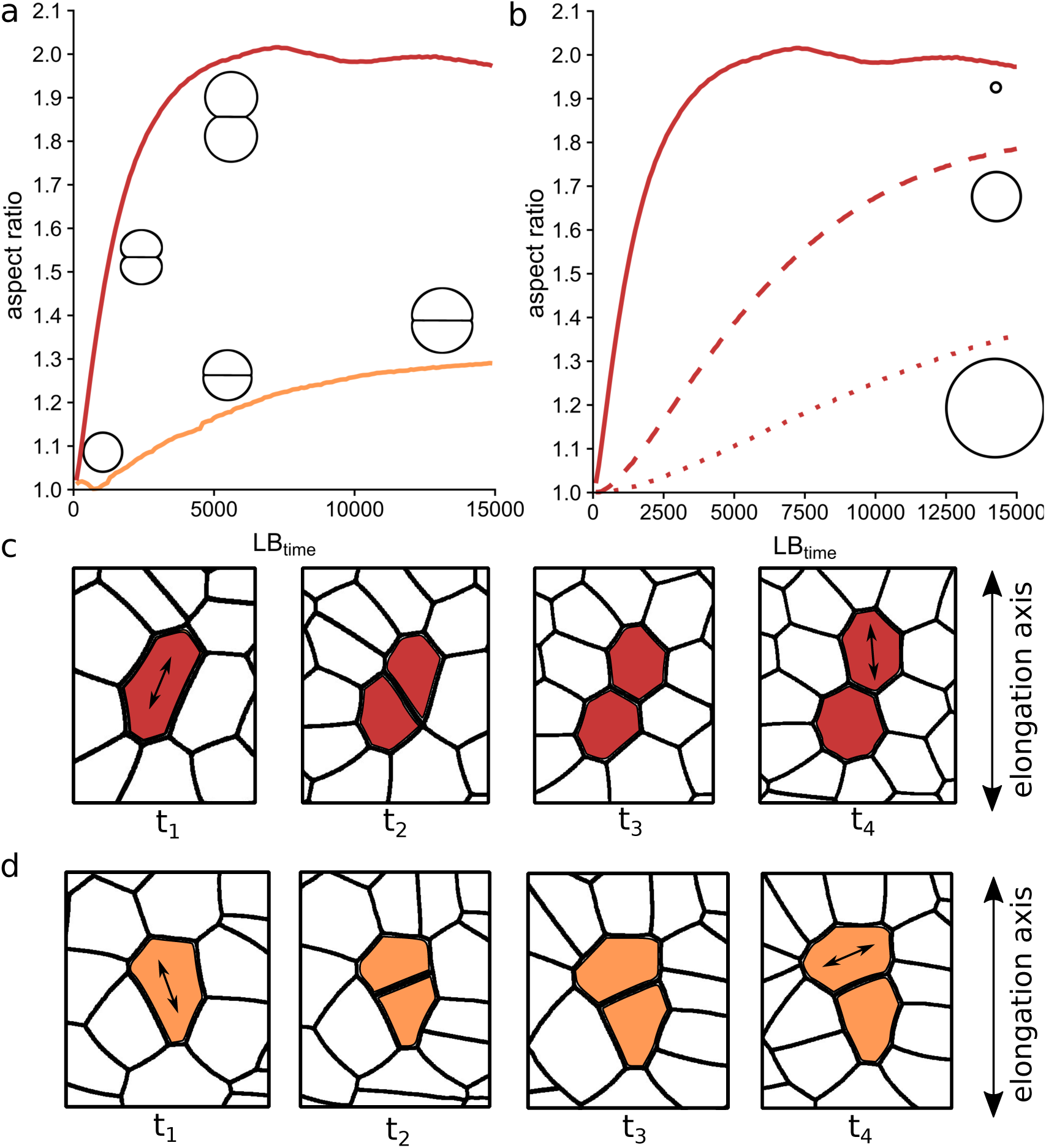
High cortical tension drives biased outgrowth by cell rounding after biased cell division. (a,b) *High cortical (membrane) tension* k_mt_ = 0.01 *(a, red) causes cell rounding after cell division, which results in a redistribution of mass in direction of cell division. This effect is weaker for lower cortical tension* k_mt_ = 0.001 *(a, orange) and for increasing cell size (b): initial cell diameter* 21LB_length_ *(solid),* 100LB_length_ *(broken),* 200LB_length_ *(dotted). The aspect ratio was determined by fitting ellipses to the outline of both cells and calculating the ratio of the main axes. The mean and standard deviation is shown for each parameter set*. (c) *In a tissue with high cortical (membrane) tension, the daughter cells round up, such that mass is distributed in direction of cell division*. (d) *In the tissue with lower membrane tension the daughter cells retain their shape after cell division and thus expand more isotropically*.

We next wondered whether the physiological cortical tension would be high enough to support elongating outgrowth. An indirect measure of the strength of the cortical tension is the fraction of hexagons in a tissue [37]. For the high cortical (membrane) tension constant that is required for biased outgrowth, the fraction of hexagons predicted by our simulations, *f*_hex_ is around 45%. Such a fraction is indeed often observed in epithelial tissues [41], and well within the measured physiological range (Figure 6a) [37]; data for the embryonic mouse epithelium is not available.

**Figure 6.**
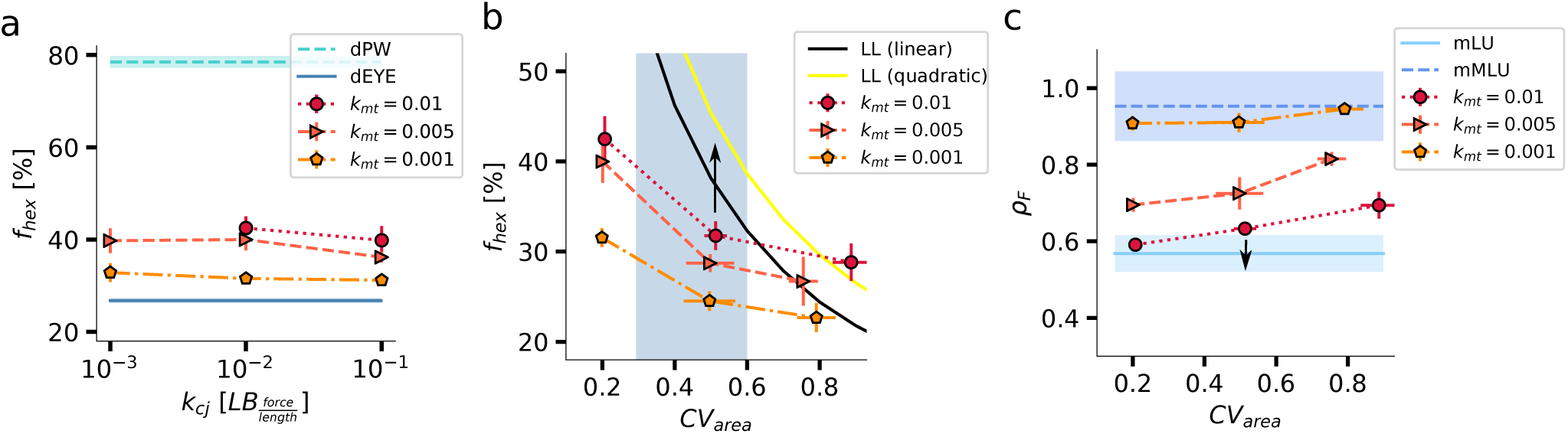
Comparison of simulated lattice organisation to measured epithelial organisation. (a) *The hexagon frequency f_hex_ depends on cortical (membrane) tension, but not on the cell-cell junction force. The hexagon frequencies in the simulations lie within the range of measured hexagon frequencies* [37]*, i.e. Drosophila pupal wing disc (dPW)* [35] *and the Drosophila eye disc (dEYE)* [36]. (b,c) *The hexagon frequency* f_hex_ (b), *and the airway shape change* ρ_F_ (c) *are affected by the area variability: for increasing* CV_area_ *the hexagon frequency is reduced and the bias in outgrowth is weaker. The simulated outgrowth ratios were compared to the measured values for wildtype* (mLU) *and mutant* (mMLU) *mouse lung epithelia* [4]*. The shaded area in* (b) *marks the range of observed* CV_area_ *values* [37]*, i.e. Drosophila pupal wing disc* (dPW) *with* CV_area_ = 0.28 [35]*, and Drosophila wing disc with gigas RNAi clones* (dMWL) *with* CV_area_ = 0.58 [37]*. The black and yellow lines represent the analytically derived relations between the fraction of hexagons and the area variability for the linear Lewis’ law (black line) and the alternative quadratic relationship (yellow line) that apply when cortical tension dominates over cell-cell adhesion* [37].

We have previously shown that the fraction of hexagons depends not only on the membrane tension constant *k*_mt_, but also on the variability of the apical cell areas [37]. Thus, most epithelial tissues follow Lewis’ law (black and yellow curves in Figure 6b [37]), and the hexagon fraction decreases as the area variability increases. The variability of the apical areas arises mainly from cell growth, cell division, and interkinetic nuclear migration. For a constant cell division threshold in our `LBIBCell` simulations, the area variability is much lower (coefficient of variation 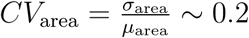) than what is observed in most tissues (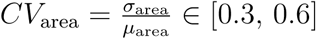, [37]); here, *µ*_area_ refers to the mean area and *σ*_area_ to its standard deviation. The area distribution can be widened by drawing the cell division threshold from a distribution rather than using a fixed value [37]. Such widening of the area distribution results in the predicted decline in the hexagon fraction [37], but most simulated tissues then lie below the predicted curves (Figure 6b), which apply when tissues seek to minimize the overall cell perimeter and thus when cortical tension dominates over cell-cell adhesion [37]. Given problems with numerical leakage [37], sufficiently high ratios between cortical tension and cell-cell adhesion strength can only be achieved in `LBIBCell` at high area CV (Figure 6b). The area CV affects the bias in outgrowth for the high membrane (cortical) tensions that are required to obtain the measured bias in outgrowth (Figure 6c). Due to the numerical limitations, the bias in outgrowth can only be reproduced for the lowest measured area CV *∼* 0.3. However, we can extrapolate that higher cortical tensions will allow us to reproduce the observed bias in outgrowth also for higher area CV values (Figure 6b,c, arrows). Such higher cortical tensions likely apply *in vivo* as all measured epithelia follow the predicted relationships between hexagon fraction are area CV that applies when cortical tension dominates over cell-cell adhesion (Figure 6b, black and yellow lines) [37]. We conclude that the observed bias in cell division is sufficient to explain the bias in outgrowth observed in the lung epithelium if cortical tension dominates over cell-cell adhesion.

### 3.2. External compressive forces can drive bias in outgrowth, but do not give rise to the measured bias in cell shape and cell division

We next investigated whether a constricting force from the surrounding tissue or ECM could explain the bias in cell shape, cell division orientation, and tube outgrowth in the lung epithelium. In our simulations, we applied compressive external forces to the growing cells and divided cells according to Hertwig’s longest axis division rule, i.e. perpendicular to their longest axis [7]. Under these conditions, we observed a bias in tissue outgrowth (Figure 7a). For comparison, we performed simulations under the same conditions without applying external forces, and we observed isotropic tissue outgrowth (Figure 7b). The velocity field along the elongation axis (y) remains linear whether or not a compressive force is applied (Figure 7c-e), thus showing that growth remains uniform. In the presence of a compressive force (*F* = 4 *×* 10*^−^*^6^ LB_force_), the velocity gradient is, however, steeper in the direction of outgrowth relative to the orthogonal direction, which is consistent with the observed biased outgrowth of *ρ_F_ ∼* 0.6 (Figure 7d).

**Figure 7.**
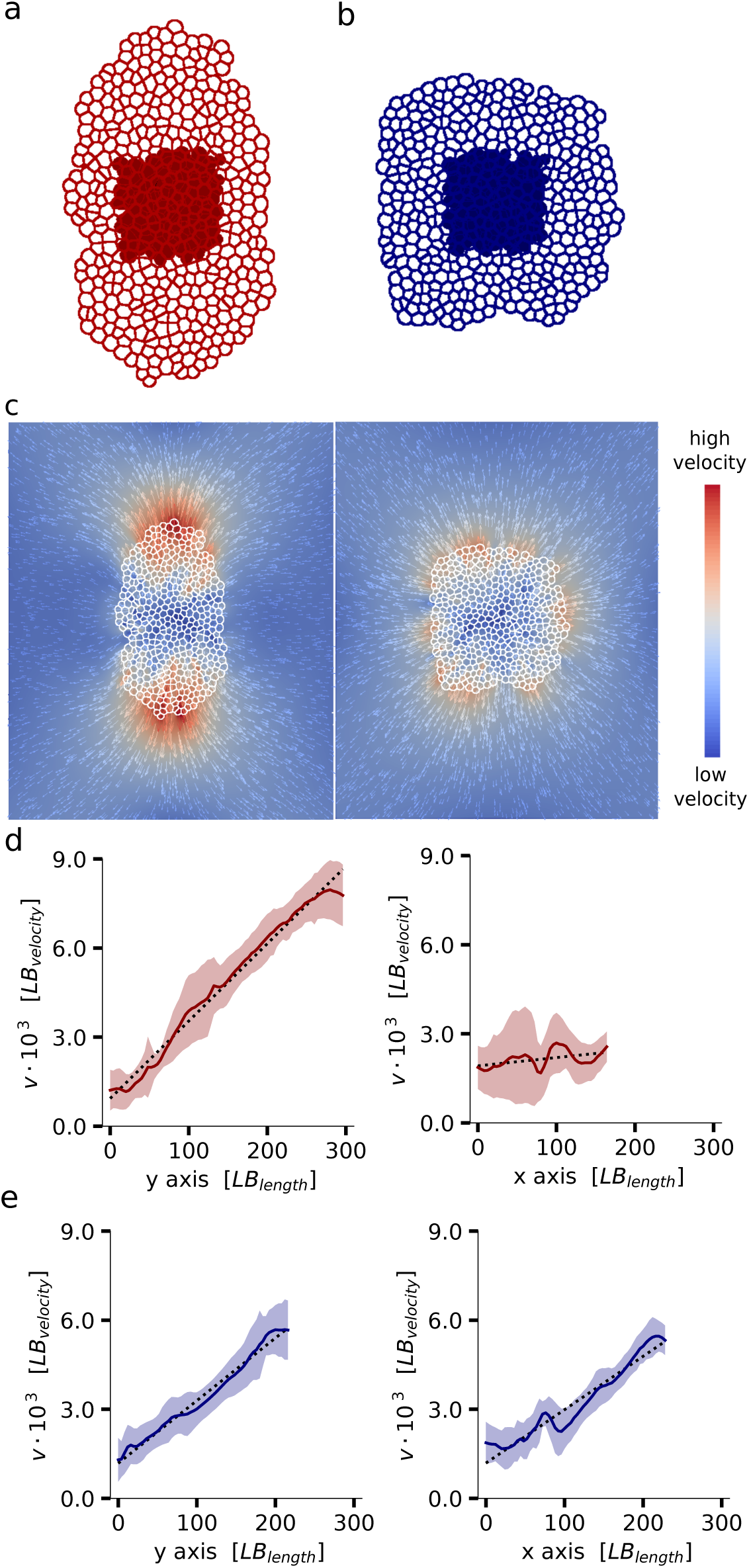
External compressive forces impact tissue outgrowth. (a,b) *Comparison of the initial tissue configuration (filled cells) with final tissue shape of the simulations with* (a) *and without* (b) *external compressive forces. The application of external forces shows anisotropic tissue expansion in the direction perpendicular to the direction of the applied forces. Simulated tissue growth without external compressive forces results in isotropic tissue expansion.* (c) *Comparison of the fluid velocity field at the end of simulations with and without external compressive forces.* (d) *Applying external compressive forces results in a large approximately linear velocity gradient along the elongation axis (y axis) and a small approximately linear velocity gradient along the x axis.* (e) *Without applying external compressive forces, a smaller approximately linear velocity gradient is observed. The gradient is similar along the x and y axis*.

We next analyzed to what extent cortical tension had an impact on tissue outgrowth and the cell division bias when applying compressive external forces. Here, we kept the cell-cell junction strength at 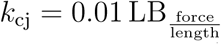, and varied only the membrane tension constant *k*_mt_ and the external forces *F*. We found that the strength of the external force determines the bias in tissue elongation, independent of the (cortical) membrane tension (Figure 8a): the higher the external force, the stronger the outgrowth perpendicular to the direction of the force, independent of the tissue’s cortical tension. The cortical tension only affected the fraction of hexagons (Figure 8b), which was slightly higher than when using the measured bias in mitotic spindle angles (Figure 6a), but still lies within the physiological range (Figure 8b). The application of external forces also affected the mitotic spindle angle distribution of a growing tissue: the fraction of biased cell divisions, i.e. the fraction of cells with a mitotic spindle angle 0*^◦^* to 10*^◦^* relative to the direction of outgrowth, increases with the applied external force (Figure 8c). However, the simulated values of cells with a biased division orientation were still much lower than the measured value in the wildtype lung and rather in the range of the measured value of the mutant lungs [4]. Plotting the bias in outgrowth, *ρ_F_*, against the fraction of biased cell divisions shows that when the measured bias in outgrowth could be reproduced, the bias in the mitotic angles was then much lower than what was observed in the lung (Figure 8d).

**Figure 8.**
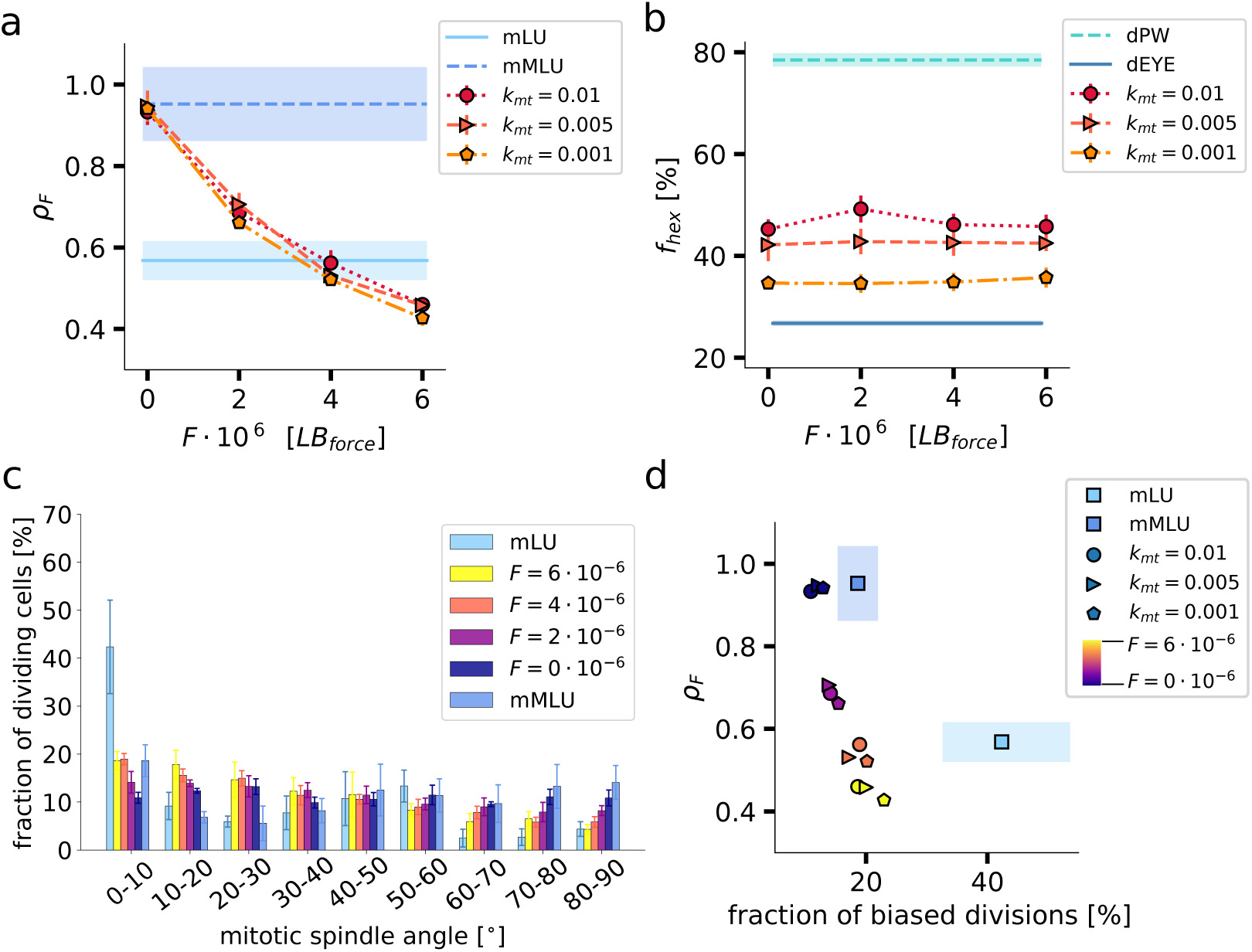
Cortical tension and external forces impact the bias of tissue outgrowth and cell division angles. (a-b) *Airway shape change ρ_F_ is decreasing for increasing external compressive force, while the hexagon frequency f*_hex_ *is not affected.* (c) *Mitotic spindle angle distributions for high k*_mt_ *show that external compressive forces increase the fraction of biased divisions.* (d) *Airway shape change ρ_F_ against fraction of biased divisions shows that for no combination of external force and membrane tension the measured values for wildtype* (mLU) *and mutant* (mMLU) *mouse lung epithelia* [4] *can be reproduced. The simulated outgrowth ratios and mitotic spindle distributions were compared to the measured values for wildtype* (mLU) *and mutant* (mMLU) *mouse lung epithelia* [4] *and hexagon frequencies were compared to measured values for epithelial tissues from Drosophila (pupal wing disc* (dPW) [35]*, eye disc* (dEYE) [36]).

In summary, the application of external compressive forces that restrain tissue expansion in one dimension led to biased tissue outgrowth and a bias in cell division orientation. However, there was no external force in our simulations, for which the measured bias in outgrowth and the measured bias in cell division for wildtype mouse lungs could both be captured.

Lastly, we tested the impact of the cell area variability for different membrane tensions in combination with constant external forces (*F* = 4 *×* 10*^−^*^6^ LB_force_). We found that for a higher cell area variability, there is a slightly smaller bias in tissue outgrowth (Figure 9a,b). A comparison to the simulations with an intrinsic bias in the mitotic spindle angle distribution (Figure 6b) showed that the impact of cell area variability on the polygon distribution of cells within a tissue was not affected by the application of external forces (Figure 9c). Corresponding to the slightly smaller bias in tissue outgrowth, the bias in the mitotic spindle angle distribution was also slightly lower for higher cell area variabilities (Figure 9d). A higher external force is thus required for larger apical area variabilities to achieve the same bias in tissue outgrowth. Overall, the external force, and to a small extent the area variability, determine biased tissue outgrowth and the mitotic spindle angle distribution, while cortical tension and area variability mainly determine the hexagon frequency of a tissue.

**Figure 9.**
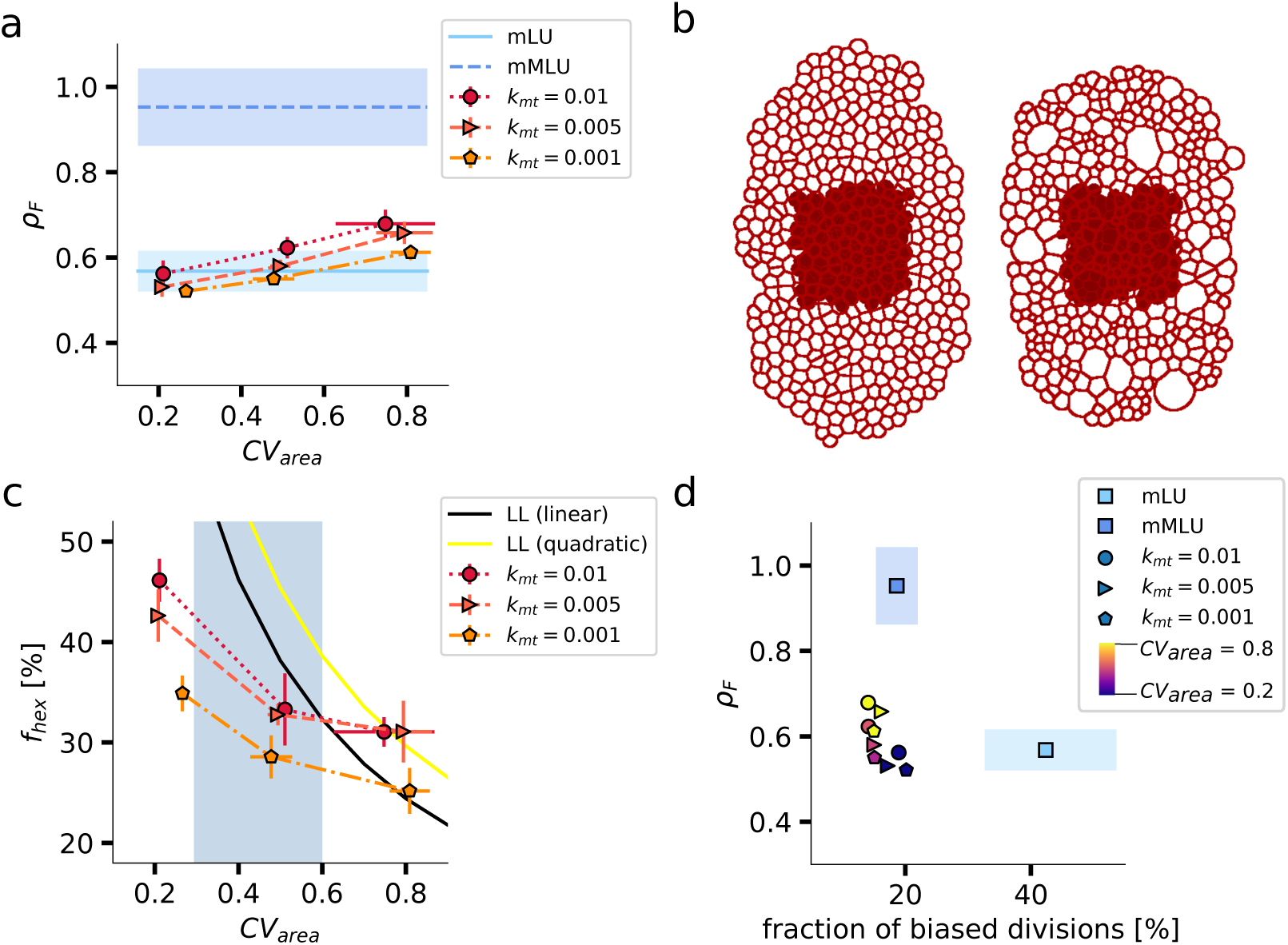
Area variability has a minor impact on the bias of tissue outgrowth and cell division angles in the presence of external forces. (a) *Airway shape change ρ_F_ increases for increasing area CV values with F* = 4 *×* 10*^−^*^6^ LB_force_. (b) *Comparing the initial tissue configuration (filled cells) with final tissue shape of the simulations, shows that for higher area CV values the anisotropic expansion is slightly lower.* (c) *Hexagon frequency f*_h_ex *decreases for increasing CV*area. (d) *Airway shape change ρF against fraction of biased cell divisions b compared to the measured values for wildtype* (mLU) *and mutant* (mMLU) *mouse lung epithelia* [4]*. The simulated outgrowth ratios were compared to the measured values for wildtype* (mLU) *and mutant* (mMLU) *mouse lung epithelia* [4] *and hexagon frequencies were compared to analytically derived relations (linear and quadratic Lewis’ law* [37]*) shown as black curve and yellow curve, respectively*.

We conclude that biased tissue outgrowth can be achieved for any cortical tension when applying compressive external forces. However, the measured bias in embryonic lung outgrowth is then achieved with a much lower bias in the cell division orientation than what is observed in experiments. This suggests that other processes generate the bias in the cell division orientation in the developing lung, and thus drive the biased outgrowth of the epithelial lung tubes during mouse development.

## 4. Discussion

We have used our cell-based tissue simulation framework `LBIBCell` to explore how biased epithelial outgrowth is achieved during mouse lung development. Experimental studies have previously linked the observed bias in outgrowth to a bias in cell shapes and cell division [4, 5], and the simulations demonstrate that this is indeed plausible as long as the cortical tension is sufficiently high. The requirement for high cortical tension is consistent with the observation that biased outgrowth is lost in mutants that express a constitutive active form of KRas (*Shhcre/*+; *KRasLSL/*+) [4]. Constitutive active KRas has previously been shown to result in a redistribution of phosphorylated myosin light chain 2 (pMLC2) inside cells of pancreatic ducts [28], which may result in reduced cortical tension on the apical side of the epithelium, where all cell divisions occur [29].

How the bias in cell shapes arises has remained unclear. We explored a possible role of constricting forces as has been proposed for the mammary gland [18] and plants [21]. While constricting forces can recapitulate the observed biased outgrowth, the resulting bias in cell shapes is much lower than what is reported for the embryonic lung epithelium. This suggests that other mechanisms drive elongating lung outgrowth. This conclusion is consistent with observations from embryonic kidneys, which show elongating outgrowth also when cultured without the surrounding mesenchyme [20]. Stretching of developing lungs in the direction of outgrowth further increases the bias in outgrowth as well as the bias in the cell division orientation [5]. Unfortunately, due to numerical limitations, the effects of stretching could not be studied using `LBIBCell`.

The biased elongation of the mammary gland has been linked to a constricting force from the myoepithelium. Based on our simulations, we would predict a low bias in cell shape in the mammary gland if this is indeed the responsible driving force. Interestingly, in the mouse embryonic neural tube, strong biased outgrowth is observed without a strong bias in cell shape [42]. The mechanism that results in biased outgrowth of the neural tube in dorsal-ventral direction is unknown [43], but our observations would support a constricting force orthogonal to the direction of biased outgrowth. As the bias in outgrowth is even higher than in the lung while the reported area CV = 0.73 is rather high in the mouse embryonic neural tube [44], a rather strong constricting force must be expected.

Going forward, it will be important to employ a combination of organ culture, live microscopy, mechanical measurements, and cell-based tissue simulations in 3D to define the mechanism that directs biased outgrowth of mammalian epithelial tubes during morphogenesis.

## Acknowledgments

This work has been supported through an SNF Sinergia grant (CRSII5 170930) to D.I. The authors thank members of the Iber group for discussions.

